# Crumbs and the Apical Spectrin Cytoskeleton Regulate R8 Cell Fate in the *Drosophila* eye

**DOI:** 10.1101/2020.10.07.329490

**Authors:** Jonathan M. Pojer, Shu Kondo, Kieran F. Harvey

**Affiliations:** Peter MacCallum Cancer Centre, 305 Grattan St, Melbourne, Victoria, Australia, 3000; Sir Peter MacCallum Department of Oncology, The University of Melbourne, Parkville, Victoria, Australia, 3010; Laboratory of Invertebrate Genetics, National Institute of Genetics, 1111 Yata, Mishima, Shizuoka, Japan; Department of Anatomy and Developmental Biology, Monash University, Clayton, Victoria, Australia, 3800

## Abstract

The Hippo pathway is an important regulator of organ growth and cell fate. In the R8 photoreceptor cells of the *Drosophila melanogaster* eye, the Hippo pathway controls the fate choice between one of two subtypes that express either the blue light-sensitive Rhodopsin 5 (Hippo inactive) or the green light-sensitive Rhodopsin 6 (Hippo active). The degree to which the mechanism of Hippo signal transduction and the proteins that mediate it are conserved in organ growth and R8 cell fate choice is currently unclear. Here, we identify Crumbs and the apical spectrin cytoskeleton as regulators of R8 cell fate. By contrast, other proteins that influence Hippo-dependent organ growth, such as the basolateral spectrin cytoskeleton and Ajuba, are dispensable for the R8 cell fate choice. Surprisingly, Crumbs promotes the Rhodopsin 5 cell fate, which is driven by Yorkie, rather than the Rhodopsin 6 cell fate, which is driven by Warts and the Hippo pathway, which contrasts with its impact on Hippo activity in organ growth. Furthermore, neither the apical spectrin cytoskeleton nor Crumbs regulate the Hippo pathway through mechanisms that have been reported to operate in growing organs. Together, these results show that only a subset of Hippo pathway proteins regulate the R8 binary cell fate decision and that aspects of Hippo pathway signalling appear to differ between growing organs and post-mitotic R8 cells.

## INTRODUCTION

Binary cell fate decisions allow for the specification of a large number of cell subtypes from a small number of precursor cells. In the nervous system, binary cell fate decisions lead to a diverse range of nearly identical cells that respond to different stimuli [1–3]. One such binary fate choice occurs in the R8 photoreceptor cells of the *Drosophila melanogaster* eye. The adult *D. melanogaster* compound eye is composed of an array of around 800 subunits, called ommatidia, each of which contains eight photoreceptor cells (R1-8). These cells are defined by a specialised subcellular compartment called the rhabdomere, which is composed of tens of thousands of microvilli that project from the cell body of each photoreceptor into the inter-rhabdomeric space at the centre of each ommatidium. The rhabdomeres of the R7 and R8 photoreceptor cells are arranged in tandem and share the same optic path, with the R7 cell positioned distally and the R8 cell positioned proximally (**Fig. 1A-A’**) [4]. Each photoreceptor cell expresses a specific rhodopsin, a photosensitive G protein-coupled receptor with a distinct spectral sensitivity [5,6]. Expression of distinct rhodopsins in different photoreceptor cells allows each cell to respond to specific wavelengths of light and prevent sensory overlap. The outer photoreceptors, R1-6, express *Rh1* and allow *D. melanogaster* to detect motion [7–9], while the inner photoreceptors, R7 and R8, express one of *Rh3*, *Rh4*, *Rh5* or *Rh6*, and are the primary cells that mediate colour vision [10].

**Figure 1.**
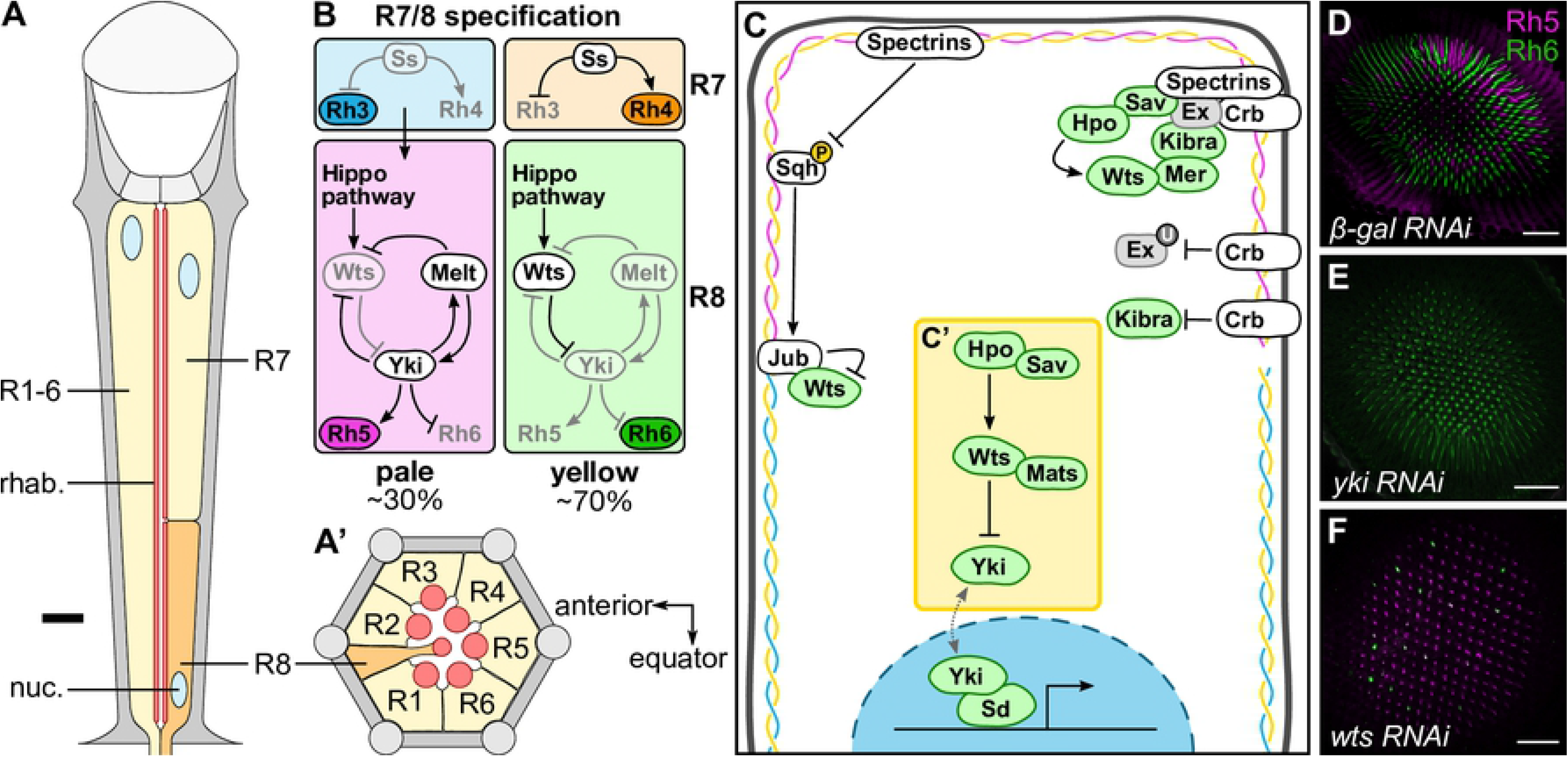
Regulation of *Drosophila melanogaster* R8 cell fate by the Hippo pathway. (**A-A’**) Schematic diagram of a *D. melanogaster* ommatidium. Yellow cells are R1-7 photoreceptor cells; orange cells are R8 photoreceptor cells; grey cells are other cells in the ommatidium. Blue circles are photoreceptor nuclei (nuc.); red lines/circles are rhabdomeres (rhab.). (**A**) Longitudinal section of an ommatidium. Note that R7 and R8 cells share the same optic path. The thick black line indicates approximately where the transverse section (**A’**) is drawn from. The distal section of the retina (towards the lens and outer surface of the eye) is to the top; the proximal section of the retina (towards the brain) is to the bottom. (**A’**) Transverse section of the proximal section of an ommatidium, showing the R8 cell. The anterior of the retina is to the left; the equator of the retina is to the bottom. (**B**) The main photoreceptor subtypes, showing R7 and R8 cell specification in each subtype. In the pale subtype, the R7 cell expresses *Rh3* (blue), signalling to the R8 cell to take on a **p**R8 cell fate through a bistable loop composed of Warts (Wts), Melted (Melt) and Yorkie (Yki) and promoting expression of *Rh5* (magenta). In the yellow subtype, the R7 cell expresses Spineless (Ss) which promotes *Rh4* (orange), while the R8 cell expresses *Rh6* (green). The subtypes are found in the specified proportions. (**C-C’**) Schematic of the Hippo pathway in tissue growth. Proteins labelled in green have already been shown to play a role in R8 cell fate; proteins labelled in grey have already been shown to not play a role in R8 cell fate; proteins labelled in white have not been studied in R8 cell fate. The spectrin cytoskeleton is shown beneath the plasma membrane, highlighting the three spectrin proteins: α-Spec (yellow), β-Spec (cyan) and Kst (magenta). The yellow box (**C’**) highlights the core kinase cassette. Crb, Crumbs; Ex, Expanded; Hpo, Hippo; Jub, Ajuba; Mats, Mob as tumour suppressor; Mer, Merlin; Sav, Salvador; Sd, Scalloped; Sqh, Spaghetti squash; Wts, Warts; Yki, Yorkie. (**D-F**) Confocal microscope images of adult *D. melanogaster* retinas stained with anti-Rh5 (magenta) and anti-Rh6 (green) antibodies. All lines were driven by *lGMR-Gal4*. Retinas expressing *β-gal RNAi* had a wild type ratio of R8 subtypes (**C**); retinas expressing *yki RNAi* had almost exclusively **p**R8 cells (**D**); retinas expressing *wts RNAi* had almost exclusively **y**R8 cells (**E**). Scale bars are 50μm.

There are different subtypes of ommatidia in the *D. melanogaster* eye, which differ based on the rhodopsins expressed in the R7 and R8 cells. The dominant subtypes are known as the ‘pale’ (**p**) and ‘yellow’ (**y**) subtypes. The **p** subtype accounts for around 30% of all ommatidia, with the short UV-sensitive *Rh3* being expressed in **p**R7 cells and the blue-sensitive *Rh5* being expressed in **p**R8 cells; the **y** subtype accounts for the remaining ~70% of ommatidia, with the long UV-sensitive *Rh4* being expressed in **y**R7 cells and the green-sensitive *Rh6* being expressed in **y**R8 cells (**Fig. 1B**). Specification of the inner photoreceptor cells is linked to ensure that the rhodopsins expressed in each subtype are always matched between R7 and R8 cells. In the late pupal retina, the transcription factor, *spineless*, is expressed stochastically in ~70% of R7 cells, inducing **y**R7 cell fate and *Rh4* expression. The remaining R7 cells take on a **p**R7 cell fate and express *Rh3* [11,12]. In these cells, the Transforming growth factor-β pathway is activated, signalling to the neighbouring R8 cell to take on a **p**R8 cell fate [13].

R8 cell fate is specified through a bistable feedback loop composed of the kinase Warts (Wts), the transcriptional coactivator Yorkie (Yki), and the Pleckstrin-homology domain protein Melted (Melt) [14,15] (**Fig. 1B**). Yki and Wts are key components of the Hippo pathway, an important regulator of organ growth and cell fate [16–18] (**Fig. 1C**). In growing organs, such as the larval imaginal discs, a kinase cassette, composed of the serine/threonine kinases, Hippo (Hpo), a sterile-20-like (Ste20) kinase [19–22], and Wts, a nuclear DBF2-related (NDR) kinase [23–25] and the scaffolding factors, Salvador (Sav) [25,26] and Mob as tumour suppressor (Mats) [27,28], inactivate the WW-domain containing transcriptional coactivator, Yki [29] (**Fig. 1C’**). Yki cannot bind to DNA itself, so must interact with transcription factors, such as the TEAD/TEF transcription factor, Scalloped (Sd), to regulate expression of target genes [30–32].

Upstream, the Hippo pathway integrates signals from surrounding cells and the extracellular matrix to regulate Yki activity [16,33–35]. Key upstream regulators of the Hippo pathway that control organ growth include the 4.1/ezrin/radixin/moesin (FERM) domain proteins, Merlin (Mer) and Expanded (Ex), and the WW-domain protein, Kibra [36–43]; the Ste20 kinase, Tao [44,45]; the polarity proteins, Crumbs (Crb), Lethal (2) giant larvae (Lgl), and the atypical cadherins Fat (Ft) and Dachsous (Ds) [42,46–52]; and mechanosensors, such as the spectrin cytoskeleton and Ajuba (Jub) [53–59] (**Fig. 1C**). These proteins localise to particular subcellular domain with many of them, such as Crb, the apical spectrin cytoskeleton, Mer, Kibra and Ex, localising to apical membrane domains and sub-apical cell junctions [43,54,60].

Many of these upstream Hippo pathway proteins also control the fate of R8 cells, which are post-mitotic. Upstream Hippo pathway proteins, such as Mer, Kibra and Lgl, converge on the core Hippo pathway kinases, Hpo and Wts in R8 cells, as in organ growth. Active Wts in **y**R8 cells prevents Yki from promoting the **p**R8 cell fate, and allows *Rh6* to be expressed [15]. Conversely, in **p**R8 cells, Wts is inactive, allowing Yki to bind to Sd and promote transcription of *Rh5*. Yki is involved in two feedback loops in R8 cells: (1) a positive feedback loop, where Yki promotes transcription of *melt*, promoting its own activation; and (2) a double-negative feedback loop, where Yki represses transcription of *wts*, thereby preventing its own repressor from acting on it [15,61] (**Fig. 1B, D-F**). This bistable feedback loop ensures that only one type of rhodopsin is expressed in each R8 cell.

Other Hippo pathway proteins, such as Ex, Ft and Ds are not required for the R8 cell fate choice [62], suggesting there are differences in how the Hippo pathway functions in different biological settings. Currently, however, we lack a complete understanding of which Hippo pathway proteins control R8 cell fate and how upstream regulators control the Hippo pathway in these cells. Here, we investigated the spectrin cytoskeleton, Crb and Jub in R8 cell fate. We identified α-Spec and Kst, components of the apical spectrin cytoskeleton, as promoters of **y**R8 cell fate and Crb as a promoter **p**R8 cell fate. By contrast, β-Spec and Jub were found to not play a role in R8 cell fate specification. Furthermore, neither the apical spectrin cytoskeleton nor Crb appear to regulate the Hippo pathway in post-mitotic R8 cells in the same manner as they do in actively growing organs.

## RESULTS

### The apical spectrin cytoskeleton promotes yR8 cell fate

To better understand Hippo signalling in R8 cells, we performed a systematic search of Hippo pathway proteins that have been implicated in organ growth but not R8 cell fate control. To assess potential roles for these proteins in R8 cell fate regulation, we genetically depleted components of the Hippo pathway in all photoreceptors using the *long Glass Multiple Reporter* (*lGMR*)*-Gal4* driver [63]. The ratio of R8 subtypes was determined by assessing the number of R8 cells that expressed Rh5 or Rh6, relative to control eyes (**Fig. 1D**, *β-gal RNAi*, approximately 30% **p**R8 cells, consistent with previous studies [14,62]). Through this screen (to be described elsewhere), we identified roles for the apicobasal polarity protein Crb and the apical spectrin cytoskeleton in the control of R8 cell fate.

The spectrin cytoskeleton is a network of large proteins that form on the intracellular surface of the plasma membrane and is widely conserved in animals [64]. In *D. melanogaster*, it is composed of tetramers of α-Spectrin (α-Spec) and one of the two β-spectrin homologues, β-Spectrin (β-Spec) or Karst (Kst, also β_Heavy_-Spectrin). These tetramers are spatially distinct in epithelial cells, with α-β tetramers localising to the basolateral membrane, and α-Kst tetramers at the apical membrane [65]. The spectrin cytoskeleton has been reported to regulate Hippo pathway activity both by responding to mechanical forces and by regulating the accumulation of upstream Hippo pathway proteins at specific plasma membrane domains [54,57]. Both spectrin cytoskeleton forms regulate the Hippo pathway in *D. melanogaster*, although differences have been reported in which of these operate in different tissues [53–56].

To investigate the role of the spectrin cytoskeleton in R8 cell fate, we depleted each spectrin gene using RNAi. Depletion of *α-Spec* (40-60% **p**R8 cells across two RNAi lines, p=0.026,<0.0001) and *kst* (43-75% **p**R8 cells across two RNAi lines, p<0.0010) in photoreceptor cells resulted in an increase in the proportion of **p**R8 cells, while depletion of *β-Spec* did not change the ratio of R8 subtypes (24-31% **p**R8 cells across three RNAi lines, p>0.055) (**Fig. 2A-D; S1A-D**). These results suggest that the apical spectrin cytoskeleton, but not the basolateral spectrin cytoskeleton, regulates R8 cell fate. This is consistent with temporally-distinct roles of the different spectrin cytoskeletons in pupal eye development, where the basolateral spectrin cytoskeleton is required for photoreceptor morphogenesis in the mid-pupal eye, while the apical spectrin cytoskeleton is required for photoreceptor morphogenesis in the late pupal eye, which coincides with when R8 subtypes are specified [66,67].

**Figure 2.**
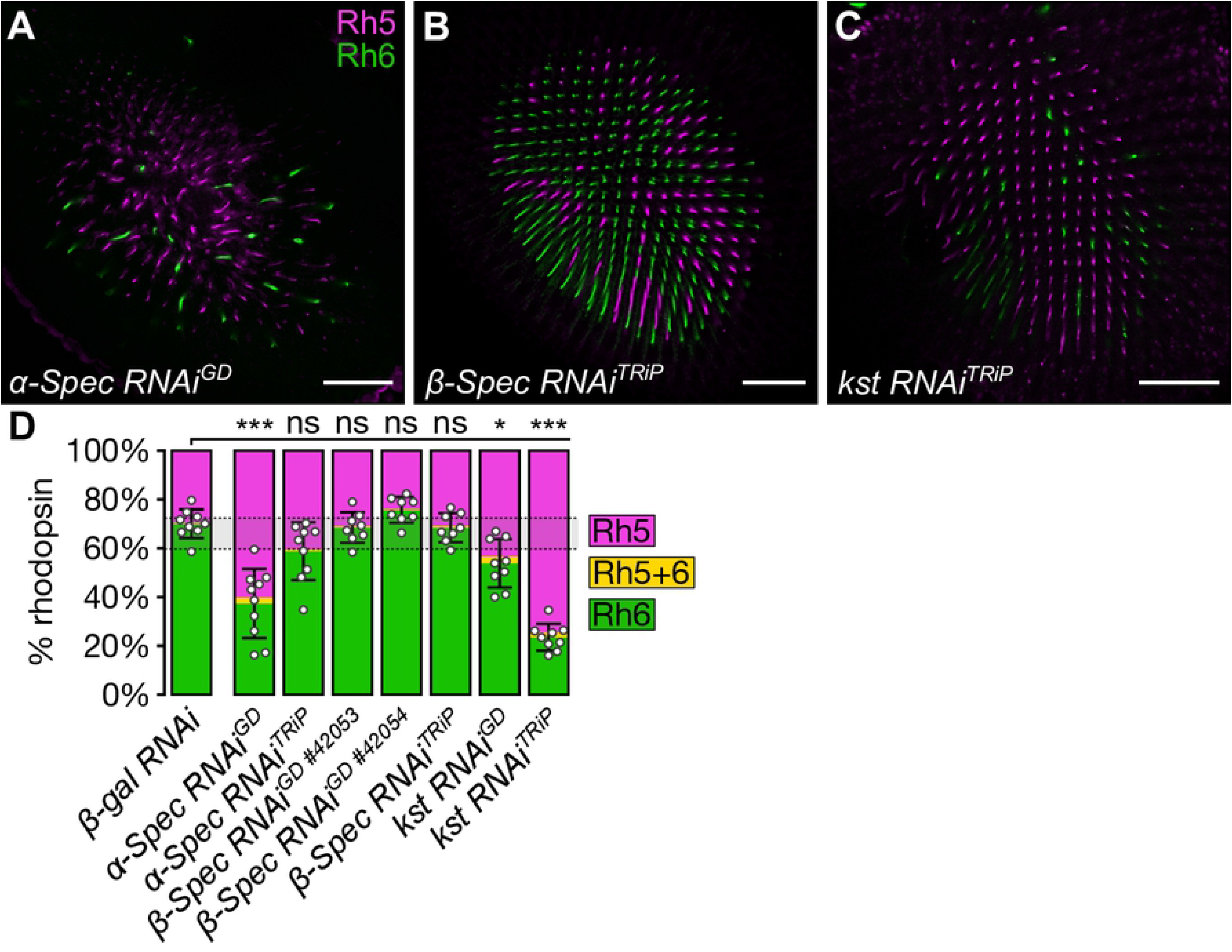
The apical spectrin cytoskeleton promotes pR8 cell fate. (**A-C**) Confocal microscope images of adult *D. melanogaster* retinas stained with anti-Rh5 (magenta) and anti-Rh6 (green) antibodies. All RNAi lines were driven by *lGMR-Gal4*. Retinas expressed *α-Spec RNAi*^*GD*^ (**A**), *β-Spec RNAi*^*TRiP*^ (**B**) and *kst RNAi*^*TRiP*^ (**C**). Scale bars are 50μm. (**D**) Proportion of R8 cells that express Rh5 (magenta), Rh6 (green), or both (yellow). The error bars represent the standard deviation of total % Rh5 (% cells expressing only Rh5 and cells co-expressing Rh5 and Rh6). Total % Rh5 was compared with two-sided, unpaired t-test; ns = not significant, * = p<0.01, *** = p<0.0001. The shaded grey region between the dotted grey lines indicates wild type Rh5:Rh6 ratio range. *β-gal RNAi* (**Fig. 1D**): n = 9 retinas, 3976 ommatidia; *α-spec RNAi*^*GD*^: n = 8, 1466; *α-spec RNAi*^*TRiP*^ (**Fig. S1A**): n = 9, 2207; *β-spec RNAi*^*GD #42053*^ (**Fig. S1B**): n = 8, 2882; *β-spec RNAi*^*GD #42054*^ (**Fig. S1C**): n = 8, 2188; *β-spec RNAi*^*TRiP*^: n = 8, 2776; *kst RNAi*^*GD*^ (**Fig. S1D**): n = 9, 3689; *kst RNAi*^*TRiP*^: n = 9, 2361.

### The subcellular localisation of α-Spectrin in R8 cells changes during late pupal eye development

Subcellular localisation is essential both for proper function of the spectrin cytoskeleton, as well as the Hippo pathway. While the two *D. melanogaster* β-Spectrin proteins, β-Spec and Kst localise at the basolateral and apical membranes, respectively, localisation of α-Spec can vary depending on which complex it forms. In the photoreceptor precursors in the larval imaginal eye disc, α-Spec localises at the apical domain, while in photoreceptors in mid-pupal eyes α-Spec localises primarily at the basolateral membrane domains and more weakly at the apical domains [66]. While the spectrins do not play an obvious role in early photoreceptor differentiation, the basal enrichment of α-Spec during the mid-pupal stage of development corresponds with an increased dependency on the basolateral spectrin cytoskeleton for morphogenesis [66].

As the Hippo pathway is important for both the specification of R8 cell fate in late pupal retinas, and the maintenance of R8 cells fate in adult eyes [62], we investigated the localisation of the apical spectrin cytoskeleton components at both stages of development. Surprisingly, 70 hours post-pupariation (hpp), when the R8 cells begin to be specified [62], α-Spec predominantly localised at the basal membrane of R8 cells, while endogenously tagged Kst (Kst-Venus) localised exclusively at the apical membrane (**Fig. 3A-A’**). This result is reminiscent of the localisation of the spectrins in photoreceptor cells during mid-pupal development, when α-Spec and β-Spec control photoreceptor morphogenesis and Kst is dispensable for this process [66]. Conversely, in adult R8 cells, which rely on the Hippo pathway to maintain their fate [62], we found that both α-Spec and Kst-Venus colocalised at the apical membrane (**Fig. 3B-B’**). Together, these results suggest that there is a switch of the dominant spectrin cytoskeleton form in R8 cells between late pupal eyes and adult eyes.

**Figure 3.**
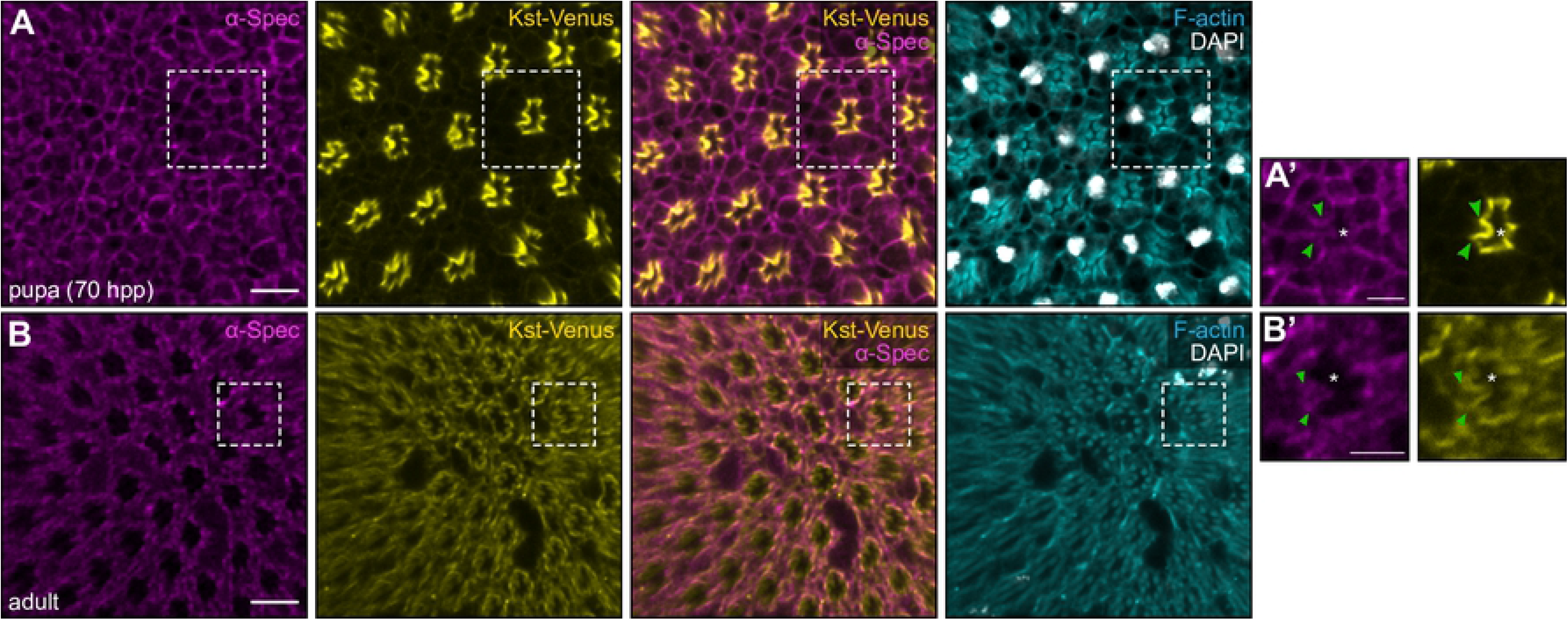
Subcellular localisation of α-Spectrin differs between late pupal and adult photoreceptor cells. (**A-B’**) Confocal microscope images of late pupal (70 hours post pupariation, hpp) and adult *D. melanogaster* retinas. Endogenously tagged Kst-Venus retinas were stained with an anti-GFP antibody to amplify the Venus signal, an anti-α-Spec antibody, DAPI (white; nuclei) and Rhodamine Phalloidin (cyan; F-actin in rhabdomeres and cell membranes). In each image, anterior is to the left. The dashed white boxes in **A** and **B** indicate the area shown in **A’** and **B’**, respectively. White asterisks indicate the rhabdomere of the ommatidium; green arrowheads indicate the adherens junctions. Scale bars are 10μm in **A** and **B**; and 5μm in **A’** and **B’**.

### The apical spectrin cytoskeleton modulates Myosin II activity but does not influence Warts localisation in R8 cells

Two models have been proposed to explain how the apical spectrin cytoskeleton influences Hippo pathway activity: (1) it regulates the phosphorylation and activation of the regulatory light chain of myosin II, Spaghetti squash (Sqh), and thereby modulates cortical tension – upon spectrin loss, cortical tension increases at adherens junctions leading to increased Jub-dependent tethering of Wts and therefore reduced Wts activity [53,56,57]; and (2) spectrins recruit core Hippo pathway proteins to the sub-apical junctions through the apicobasal polarity protein Crb – upon spectrin loss, Hippo activation complexes are depleted and Wts activity is reduced [54] (**Fig. 1C**). Consistent with published studies [57], upon depletion of either *α-Spec* or *kst*, phosphorylated Sqh (pSqh) was increased in photoreceptor cells (**Fig. 4A-D**). To investigate the effect of this increase of pSqh on the Hippo pathway, we assessed the localisation of a Jub-GFP transgene [68]. While Jub-GFP colocalised with E-Cadherin at adherens junctions in larval eye imaginal discs, we could not detect Jub-GFP expression in adult photoreceptor cells (**Fig. 4E-F**). To further interrogate this, we investigated a potential role for *jub* in R8 cell fate and found that expression of *jub* RNAi lines did not alter the R8 subtype ratio (21-30% **p**R8 cells across two RNAi lines, p>0.023) (**Fig. 4G-H**; **S1E**). These results suggest that while the apical spectrin cytoskeleton regulates phosphorylation of Sqh in R8 cells, this does not mediate its impact on the Hippo pathway, given that we found no role for *jub* in R8 cell fate and could not detect its expression in these cells.

**Figure 4.**
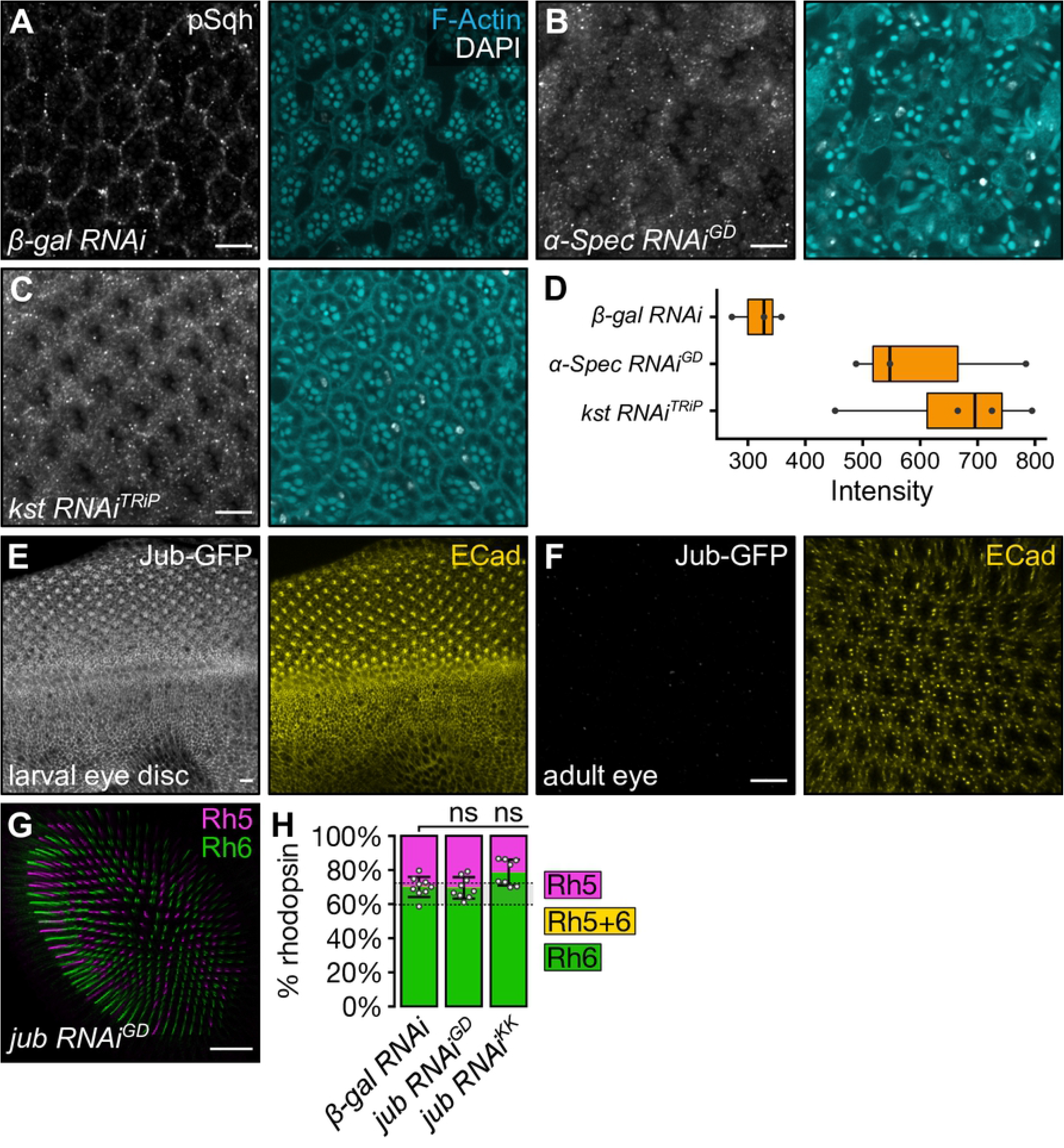
The apical spectrin cytoskeleton modulated phosphorylation of Spaghetti squash. (**A-C**) Confocal microscope images of adult *D. melanogaster* stained with anti-pSqh (grey) antibody, DAPI (white) and Phalloidin (F-actin, cyan). All RNAi lines were driven by *lGMR-Gal4*. Retinas expressed *β-gal RNAi* (**A**), *α-Spec RNAi*^*GD*^ (**B**), and *kst RNAi*^*TRiP*^ (**C**). Scale bars are 10μm. (**D**) Boxplot showing intensity of pSqh in (**A-C**). (**E-F**) Confocal microscope images of *Jub-GFP D. melanogaster* in a larval eye imaginal disc (**E**) and an adult eye (**F**). Tissues were stained with anti-GFP (grey) and anti-ECad (yellow) antibodies. Scale bars are 10μm. (**G**) Confocal microscope images of adult *D. melanogaster* retina stained with anti-Rh5 (magenta) and anti-Rh6 (green) antibodies. Expression of *jub RNAi*^*GD*^ was driven by *lGMR-Gal4*. Scale bar is 20μm. (**H**) Proportion of R8 cells that express Rh5 (magenta), Rh6 (green), or both (yellow). The error bars represent the standard deviation of total % Rh5 (% cells expressing only Rh5 and cells co-expressing Rh5 and Rh6). Total % Rh5 was compared with two-sided, unpaired t-test; ns = not significant. The shaded grey region between the dotted grey lines indicates wild type Rh5:Rh6 ratio range. *β-gal RNAi* (**Fig. 1D**): n = 9 retinas, 3976 ommatidia; *jub RNAi*^*GD*^: n = 9, 3660; *jub RNAi*^*KK*^ (**Fig. S1E**): n = 8, 3048.

### Crumbs promotes pR8 cell fate

An alternative mechanism by which the apical spectrin cytoskeleton has been proposed to regulate the Hippo pathway is by forming a complex with Crb at sub-apical junctions [54]. Kst and Crb physically interact in a number of *D. melanogaster* tissues, including embryos and pupal photoreceptors, where they promote correct apical domain formation [66,67,69]. In larval wing imaginal discs, Crb and Kst colocalise at the sub-apical junction and have been reported to promote accumulation of several Hippo pathway proteins at these junctions, including Ex, Mer, Kib, Hpo and Wts, leading to Hippo pathway activation and suppression of Yki-mediated tissue growth [54]. As Crb and Kst colocalise in pupal and adult photoreceptor cells at the stalk membrane (the apical membrane below the rhabdomere [66,67]), we hypothesised that they also recruit Hippo pathway proteins to the stalk membrane and promote its activation in this membrane domain. To investigate a role for Crb in R8 cell fate, we used *eyFlp/FRT* site-specific recombination [70] to generate clones of tissue harbouring the *crb* null allele, *crb*^*11A22*^ [71] (**Fig. 5A-B, D**; **S2A, C**). *crb*^*11A22*^ clones displayed two distinct phenotypes that distinguished them from wild type clones. First, *crb*^*11A22*^ clones had a reduction in the proportion of **p**R8 cells by around 3.6 times when compared with neighbouring wild type clones (**Fig. 5A, D**; **S2C**). This was surprising, as it suggests that Crb actually promotes **p**R8 cell fate, while the apical spectrin cytoskeleton and other Hippo pathway proteins with which Crb has been reported to function within the context of epithelial tissue growth, such as Wts, Hpo and Sav, promote the opposing **y**R8 cell fate. Second, the rhabdomeres of photoreceptors in *crb*^*11A22*^ clones were shortened, suggesting a fault in photoreceptor morphogenesis (**Fig. S2A**), a phenotype that has been previously described as a failure of rhabdomere elongation in pupal development [72].

**Figure 5.**
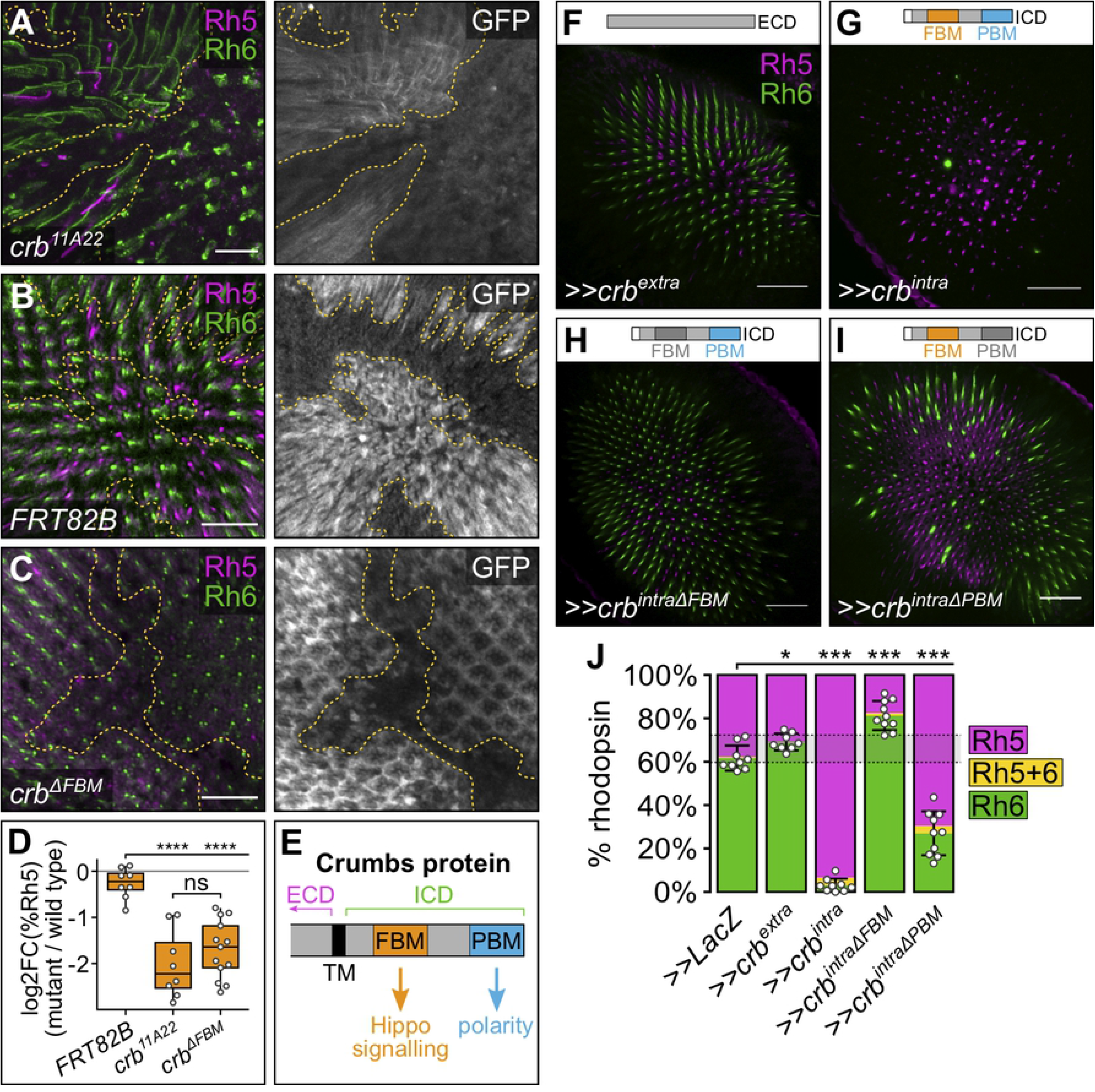
Crumbs regulates R8 cell fate through its FERM-binding motif. (**A-C**) Confocal microscope images of adult *D. melanogaster* retinas stained with anti-GFP (grey), anti-Rh5 (magenta) and anti-Rh6 (green) antibodies. GFP-negative clones were mutant for *crb*^*11A22*^ (**A**), *FRT82B* (negative control) (**B**) and *crb*^*ΔFBM*^.*HA* (**C**). Panel (**A**) is a maximum projection as rhodopsins localise in different planes in wild type and mutant clones. Scale bars are 20μm. (**D**) Log2 value of the ratio of total % Rh5 (% cells expressing only Rh5 and cells co-expressing Rh5 and Rh6) between mutant and wild type clones from the same tissue. Genotypes were compared with an ANOVA; ns = not significant; **** = p<0.0001. *FRT82B*: n = 8 retinas, 4065 ommatidia; *crb*^*11A22*^: n = 8, 1394; *crb*^*ΔFBM*^.*HA*: n = 10, 3851. (**E**) Schematic illustration of the intracellular domain of the Crb protein. ECD, extracellular domain; ICD, intracellular domain; TM, transmembrane domain; FBM, FERM-binding motif; PBM, PDZ-binding motif. (**F-I**) Confocal microscope images of adult *D. melanogaster* retinas stained with anti-Rh5 (magenta) and anti-Rh6 (green) antibodies. All lines were driven by *lGMR-Gal4*. Retinas expressed *crb*^*extra*^ (**F**), *crb*^*intra*^ (**G**), *crb*^*intraΔFBM*^ (**H**) and *crb*^*intraΔPBM*^ (**I**). Schematic illustrations in the top right corner indicate the transgenes expressed in each experiment; motifs in dark grey indicate mutated motifs in the transgene. Scale bars are 20μm. (**J**) Proportion of R8 cells that express Rh5 (magenta), Rh6 (green), or both (yellow). The error bars represent the standard deviation of total % Rh5 (% cells expressing only Rh5 and cells co-expressing Rh5 and Rh6). Total % Rh5 was compared with two-sided, unpaired t-test; * = p<0.01, *** = p<0.0001. The shaded grey region between the dotted grey lines indicates wild type Rh5:Rh6 ratio range. ≫*LacZ*: n = 9 retinas, 3211 ommatidia; ≫*crb*^*extra*^: n = 8, 2153; ≫*crb*^*intra*^: n = 9, 2381; ≫*crb*^*intraΔFBM*^: n = 10, 3351; ≫*crb*^*intraΔPBM*^: n = 10, 4041.

### Crumbs regulates R8 cell fate through its FERM-binding motif

A question that arose from these observations was whether these two phenotypes – the change in R8 subtype ratio and the disruption of rhabdomere morphogenesis – were linked. Crb is a transmembrane protein composed of a long extracellular domain, a transmembrane domain, and a short intracellular domain. The extracellular domain is essential for Crb apical enrichment and stabilisation, and cell aggregation by mediating Crb-Crb interactions between neighbouring cells [73,74]. The intracellular domain contains two defined motifs – a juxtamembrane FERM-binding motif (FBM) and a C-terminal PDZ (PSD-95/Dlg/ZO-1)-binding motif (PBM) (**Fig. 5E**). The Crb FBM can bind to FERM domains in proteins such as Ex, to regulate Hippo pathway activity and tissue growth [42,46–48], while the PBM recruits members of the Crumbs complex (Stardust, Patj and Lin-7) to promote and maintain apicobasal polarity and epithelial integrity [75]. To investigate the role of the Crb FBM in R8 cell fate, we generated clones for *crb^ΔFBM^*, an allele with mutations to three key residues in the Crb FBM [42] (**Fig. 5C-D**, **S2C**). Notably, in *crb*^*ΔFBM*^ R8 cells, rhodopsin localisation extended the whole length of the R8 cell, as in neighbouring wild type R8 cells, indicating that these mutations do not affect rhabdomere morphogenesis, as seen in *crb*^*11A22*^ cells. Furthermore, *crb*^*ΔFBM*^ clones showed on average a 3.3-fold decrease in the percentage of **p**R8 cells compared to neighbouring wild type clones (**Fig. 5C**, **S2C**). The magnitude of change in R8 cell ratio was very similar to that observed in *crb*^*11A22*^ clones (p=0.981) (**Fig. 5D**) and indicates that Crb promotes **y**R8 cell fate through its FBM.

To investigate whether *crb* overexpression is sufficient to perturb R8 cell fate choice, we misexpressed transgenes composed of only the *crb* extracellular (*crb*^*extra*^) or intracellular (*crb*^*intra*^) domains in all photoreceptor cells (**Fig. 5F-G, J**). In *lGMR*>*crb*^*extra*^ eyes, there was a mild, but statistically significant, decrease in the proportion of **p**R8 cells (approximately 31% **p**R8 cells, p=0.0074) compared to the *lGMR*>*LacZ* control (approximately 38% **p**R8 cells), though the average proportion of Rh5-positive **p**R8 cells was still within the wild type range (**Fig. 5F, J**). Strikingly however, *lGMR*>*crb*^*intra*^ eyes, had a strong increase in the proportion of **p**R8 cells (around 93% **p**R8 cells, p<0.0001) (**Fig. 5G, J**). Therefore, in R8 cells, mutating *crb* or misexpressing *crb*^*intra*^ led to opposing phenotypes – i.e. a decrease or increase in the proportion of **p**R8 cells, respectively, which phenocopies genetic analysis of the Yki transcription coactivator in R8 cells [15]. Interestingly, this role for Crb in R8 cells contrasts with its role in larval eye and wing imaginal disc, where mutation of *crb* and misexpression of *crb*^*intra*^ both cause Yki hyperactivation and tissue overgrowth [42,46–48]. These data suggest that there are important differences in how Crb signals to the Hippo pathway in post-mitotic R8 cells and in growing organs.

As mutation of the *crb* FBM was sufficient to alter the ratio of R8 subtypes to the same extent as a *crb* null allele, we hypothesised that mutating the FBM in the *crb*^*intra*^ transgene (*crb*^*intraΔFBM*^) would abolish the effects of *crb*^*intra*^ misexpression on R8 cell fate. Indeed, *lGMR*>*crb*^*intraΔFBM*^ failed to shift the balance of R8 cells to the **p**R8 fate and, in fact, slightly shifted it to the **y**R8 fate (approximately 18% **p**R8 cells, p<0.0001) (**Fig. 5H, J**). This suggested that another part of the Crb intracellular domain plays a minor role in R8 cell fate control, with a likely candidate being the PBM, which is essential to Crb’s role in photoreceptor morphogenesis [72]. To investigate this, we misexpressed a *crb*^*intra*^ transgene with a mutated PBM (*crb*^*intraΔPBM*^) in photoreceptor cells. *lGMR*>*crb*^*intraΔPBM*^ eyes had an increased proportion of **p**R8 cells compared to *lGMR*>*LacZ* (around 70% **p**R8 cells, p<0.0001), though not to the same extent as in *lGMR*>*crb*^*intra*^ eyes (p<0.0001) (**Fig. 5I-J**). Collectively, this suggests that while the Crb FBM plays a major role in promoting **p**R8 cell fate, the PBM plays a minor role in promoting **y**R8 cell fate.

### Crumbs regulates R8 cell fate independent of Kibra

In *D. melanogaster* larval imaginal discs, Crb can regulate the Hippo pathway by interacting with different upstream Hippo pathway proteins (**Fig. 1C**). Crb directly interacts with Ex, both recruiting it to the apical membrane to promote activation of the pathway [42,46,48,54,60,76,77] and promoting ubiquitin-mediated degradation of Ex [78,79]. Crb also represses Kibra by sequestering it at sub-apical junctions. In the absence of Crb, Kibra localises at the medial apical cortex and recruits Mer, Sav and Wts to this membrane domain, thus activating the Hippo pathway core cassette [43]. As Ex does not regulate R8 cell fate [62], we hypothesised that Crb might regulate the Hippo pathway in R8 cells by repressing Kibra. To investigate the relationship between Crb and Kibra in R8 cells, we generated both *kibra*^*4*^ *crb*^*11A22*^ and *kibra*^*4*^ *crb*^*ΔFBM*^ double mutant clones. Mutant clones for *kibra*^*4*^, a null *kibra* allele [37], had a dramatic expansion (10.2 times higher) in the proportion of **p**R8 cells relative to wild type clones (**Fig. S2B-C**) [62]. Similarly, both *kibra*^*4*^ *crb*^*11A22*^ and *kibra*^*4*^ *crb*^*ΔFBM*^ clones also had a greatly increased proportion of **p**R8 cells compared to wild type cells from the same tissue (15 and 26 times higher, respectively) (**Fig. 6A-C, S2C**), suggesting that Kibra acts downstream of, or in parallel to, Crb in R8 cells.

**Figure 6.**
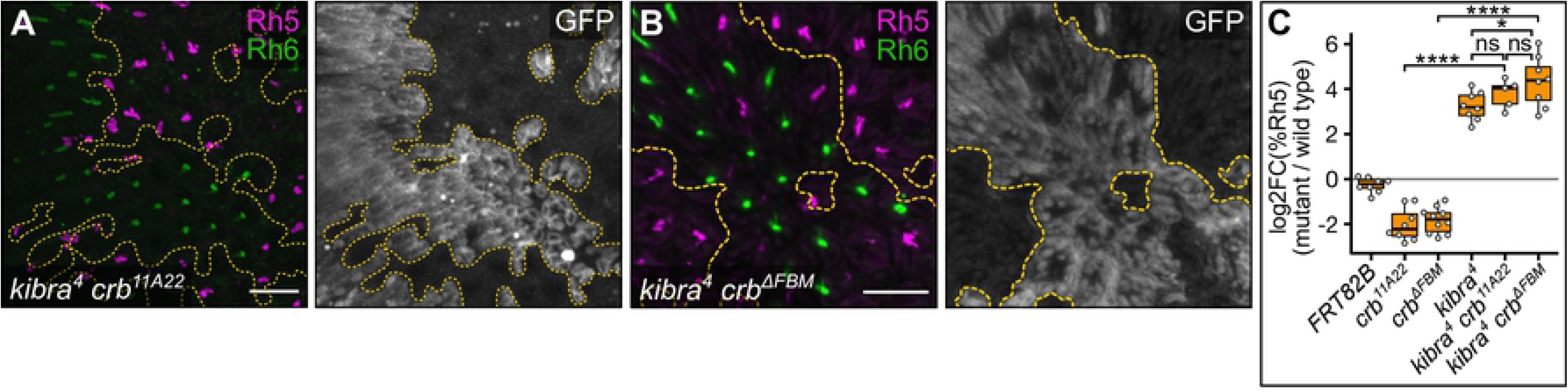
Crumbs does not affect the subcellular localisation of Kibra in R8 cells. (**A-B**) Confocal microscope images of adult *D. melanogaster* retinas stained with anti-GFP (grey), anti-Rh5 (magenta) and anti-Rh6 (green) antibodies. GFP-negative clones were mutant for *kibra*^*4*^ *crb*^*11A22*^ (**A**), and *kibra*^*4*^ *crb*^*ΔFBM*^ (**B**). Panel (**A**) is a maximum projection as rhodopsins localise in different planes in wild type and mutant clones. Scale bars are 20μm. (**C**) Log2 value of the ratio of total % Rh5 (% cells expressing only Rh5 and cells co-expressing Rh5 and Rh6) between mutant and wild type clones from the same tissue. Genotypes were compared with an ANOVA; ns = not significant; *** = p<0.001. *FRT82B* (**Fig. 4B**): n = 8 retinas, 4065 ommatidia; *crb*^*11A22*^ (**Fig. 4A**): n = 8, 1394; *crb*^*ΔFBM*^.*HA* (**Fig. 4C**): n = 10, 3851; *kibra*^*4*^: n = 8, 2776; *kibra*^*4*^ *crb*^*11A22*^: n = 5, 1174; *kibra*^*4*^ *crb*^*ΔFBM*^: n = 8, 2479. (**D-E**) Confocal microscope images of adult *D. melanogaster* retinas stained with anti-GFP (grey) antibody and anti-HA antibody (yellow) or Phalloidin (F-actin, cyan). GFP-positive clones expressed *kibra-Venus*, in wild-type cells (**D)** or cells harbouring the *crb*^*ΔFBM*^.*HA* allele (**E**). Scale bars are 5μm. (**F-H**) Confocal microscope images of *kibra-Venus* adult *Drosophila* retinas stained with anti-GFP (grey) antibody. All lines were driven by *lGMR-Gal4*. Retinas expressed *crb*^*intra*^ (**F**), *crb*^*intraΔFBM*^ (**G**) and *crb*^*intraΔPBM*^ (**H**). Scale bars are 5μm. (**I-J**) Confocal microscope images of *Mer-Venus* adult *D. melanogaster* retinas stained with anti-GFP (grey) and anti-Rh6 (green) antibodies. All lines were driven by *lGMR-Gal4*. Retinas expressed no transgene (**I**) and *kibra* (**J**). Yellow stars indicate the R8 rhabdomere; green arrows indicate the stalk of the R8 cell. Scale bars are 5μm.

To investigate this further, we assessed whether Crb regulates Kibra subcellular localisation in R8 cells. To do this, we generated an endogenously tagged Kibra-Venus *D. melanogaster* strain using CRISPR-Cas9 genome editing (to be described elsewhere). In wild-type adult R8 photoreceptor cells, Kibra-Venus was only weakly expressed and visible throughout the cytoplasm (**Fig. 6D**). We predicted that in the absence of *crb*, Kibra-Venus would relocalise from the cytoplasm to the rhabdomere, a potentially analogous membrane domain to the medial apical cortex in larval imaginal discs. However, the subcellular localisation of Kibra-Venus was unaltered in *crb*^*ΔFBM*^ clones (**Fig. 6E**). Similarly, misexpression of the *crb*^*intra*^ transgene in photoreceptor cells did not alter Kibra-Venus localisation, nor did misexpression of either the *crb*^*intraΔFBM*^ or *crb*^*intraΔPBM*^ transgenes (**Fig. 6F-H**). Collectively, these data suggest that Crb does not obviously regulate the Hippo pathway by controlling Kibra subcellular localisation in R8 cells, as it does in growing wing imaginal discs.

### Kibra and Merlin do not regulate the Hippo pathway at the rhabdomere

In larval imaginal wing discs, Kibra recruits Hippo pathway components, notably Mer, Sav, Hpo and Wts, to the medial apical cortex to promote Hippo pathway activation [43]. We predicted that if Kibra regulates the Hippo pathway at the medial apical cortex in R8 cells as it does in larval imaginal discs, Kibra overexpression would cause Mer to accumulate at the rhabdomere. To visualise Mer in R8 cells, we generated an endogenously tagged Mer-Venus *D. melanogaster* strain using CRISPR-Cas9 genome editing (to be described elsewhere). We found that in both control and *kibra*-overexpressing R8 cells, Mer-Venus predominantly localised at the stalk membrane (**Fig. 6I-J**). This suggests that in R8 cells, the Hippo pathway is not activated at membrane domains that are analogous to the medial apical cortex of wing imaginal disc cells.

## DISCUSSION

The Hippo pathway is a complex signalling network that integrates multiple signals to control organ growth and cell fate decisions, including the binary fate choice of R8 photoreceptors in the *D. melanogaster* eye [14]. The proteins that take part in Hippo pathway signal transduction in organ growth are better understood than those in cell fate. Here, we have identified the apical spectrin cytoskeleton proteins α-Spec and Kst, and the apicobasal polarity protein Crb, as important regulators of the R8 cell fate choice. By contrast, we found that neither β-Spec nor Jub, which operate in the Hippo pathway in tissues such as the ovary and the imaginal discs, do not regulate R8 cell fate. Therefore, these studies provide new information on R8 cell fate specification and how the Hippo pathway mediates signal transduction in different biological settings.

Interestingly, our study suggests that Crb and the apical spectrin cytoskeleton each transduce signals to the Hippo pathway via distinct modes in organ growth as opposed to R8 cell fate. While both mutation and overexpression of *crb* in growing wing and eye imaginal discs leads to Yki hyperactivity and tissue overgrowth [42,46–48], loss of *crb* in R8 cells led to a decrease in **p**R8 cells – synonymous with reduced Yki activity – while *crb* overexpression caused an increase in **p**R8 cells – a phenotype linked to increased Yki activity. In growing larval imaginal discs, Crb has been reported to have opposing influences on Hippo pathway activity through three mechanisms: (1) it recruits Ex to the sub-apical junctions, leading to Hippo pathway activation; (2) it promotes the ubiquitination and degradation of Ex, resulting in suppression of Hippo pathway activity; and (3) it sequesters Kibra at sub-apical junctions, limiting activation of the Hippo pathway at the medial apical cortex (**Fig. 7A**) [42,43,46–48,78]. In growing larval imaginal discs, Ex’s role as an activator of the Hippo pathway must dominate over the other mechanisms, as Crb loss impedes Hippo pathway activity [42,46–48]. By contrast, in R8 cells, Crb appears to primarily mediate a Hippo-inhibitory signal that results in Yki activation. This apparent change in Crb signalling to the Hippo pathway could be explained by the low expression of Ex in pupal and adult eyes [80] and the dispensability of *ex* for R8 cell fate [62]. Accordingly, only the Kibra-antagonism function of Crb might operate in R8 cells. Consistent with this, we found that *kibra* loss was completely epistatic to *crb* loss, with respect to R8 cell fate choice. However, unlike in growing imaginal discs [43], we found no evidence that Crb influences the subcellular localisation of Kibra in R8 cells, although these studies were technically challenging because of the very low expression of Kibra. Our genetic analysis of *crb* in R8 cells suggests a fourth mechanism by which Crb signals to the Hippo pathway via an unidentified FERM-domain protein that suppresses Hippo pathway activity (**Fig. 7B**). Candidate FERM-domain proteins that might regulate the Hippo pathway in conjunction with Crb include Moesin (Moe) and Yurt (Yrt), which have both been shown to physically interact with Crb [69,81,82]. While neither Moe nor Yrt have been directly associated with Hippo signalling, Moe associates with both Crb and Kst [69] and plays a role in photoreceptor morphogenesis [83], while Yrt is a negative regulator of Crb in photoreceptor development [81,82]. This putative Crb-dependent regulatory mechanism of the Hippo pathway might be R8-specific or also operate more broadly, for example in growing larval imaginal discs.

**Figure 7.**
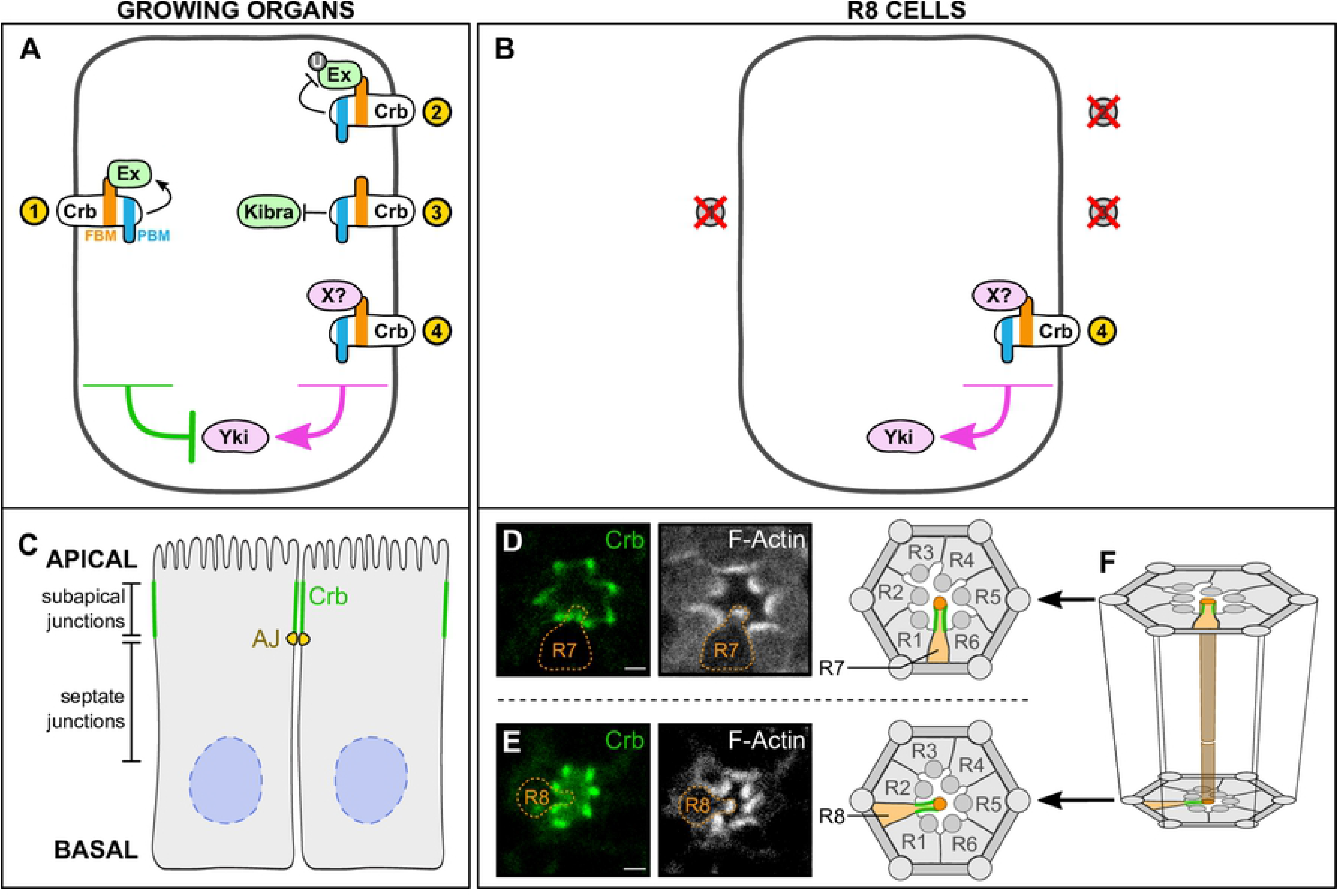
Model of the Hippo pathway in R8 cells. (**A-B**) Schematic diagram of the role of Crb in the Hippo pathway in growing organs (**A**) and R8 cells (**B**). Proteins and arrows in magenta promote organ growth or **p**R8 cell fate; proteins and arrows in green suppress organ growth or promote **y**R8 cell fate. Crb, Crumbs; Ex, Expanded; FBM, FERM-binding motif; PBM, PDZ-binding motif; Yki, Yorkie. (**C-F**) Localisation of Crb in epithelial cells and R8 photoreceptor cells. (**C**) Schematic diagram of epithelial cells, showing Crb localisation (green) at the sub-apical junctions. Adherens junctions (AJ) are shown in yellow; nuclei are shown in blue. (**D-E**) Confocal microscope images of a *Crb-GFP* pupal ommatidium stained with Phalloidin (F-Actin). R7 and R8 cells are outlined in orange. Scale bars are 2μm. (**F**) Schematic diagram of an ommatidium, showing R7 and R8 planes from (**D**) and (**E**). The brown tube at the centre of the diagram indicates the optic path shared by the rhabdomeres of the R7 and R8 cells. Crb is shown in green; R7 and R8 cells are shown in orange.

Another important consideration regarding the role of Crb in R8 cell fate is whether it conveys signals that are mediated by its extracellular domain. In larval imaginal discs, Crb engages in homophilic interactions between apical junctions of neighbouring cells and this has been hypothesised to be important for its ability to control Hippo signalling and cell competition [84]. The Crb extracellular domain is also important for stability of the Crb protein and the maintenance of apicobasal polarity [73]. The role of the Crb extracellular domain in R8 cell fate is currently unclear although its misexpression did not influence R8 cell fate choice. R8 fate is induced by the neighbouring R7 cells [85–87] and is conveyed by parallel Activin and BMP signalling, however how Hippo pathway activity is influenced in R8 cells has not yet been elucidated [88]. One possibility is that Crb engages in homophilic interactions and thereby signals from R7 to R8 cells, although based on its subcellular localization in these cells, we think this is unlikely. In photoreceptor cells, Crb localises at the membrane of the stalk, the apical subcellular compartment of photoreceptor cells located basally of the rhabdomere [72,89], analogous to its localisation at the sub-apical junctions in imaginal disc epithelial cells [75] (**Fig. 7C-F**). However, while Crb is directly apposed in neighbouring imaginal disc epithelial cells and allows for interactions between Crb extracellular domains of adjacent cells (**Fig. 7C**), the stalks of the R7 and R8 cells do not obviously overlap (**Fig. 7C-F**). Therefore, Crb is unlikely to signal between neighbouring R7 and R8 cells via homophilic interactions between the Crb extracellular domain.

Another apparent difference in Hippo signalling between growing organs and R8 cells relates to the apical spectrin cytoskeleton. As described above, in growing larval imaginal discs, the apical spectrin cytoskeleton has been proposed to regulate the Hippo pathway by two modes: (1) by binding to both Crb and Ex, which recruit the core kinase cassette to sub-apical junctions to be activated [54]; and (2) by influencing cytoskeletal tension and, thereby, Jub-dependent suppression of Wts at adherens junctions [53,57]. Here, we found that depletion of Crb and the apical spectrin cytoskeleton components, α-Spec and Kst, have an opposing impact on R8 cell fate. This suggests that their regulatory roles are decoupled in the context of R8 cell fate, which is particularly surprising since Crb and the apical spectrin cytoskeleton co-ordinately regulate photoreceptor morphogenesis during pupal development [67]. Additionally, we showed that depletion of the apical spectrin cytoskeleton leads to an increase of pSqh, as in the pupal eye [53]. However, we were unable to detect expression of Jub in adult photoreceptors and found no role for Jub in R8 cell fate choice. As such, the mechanism by which the apical spectrin cytoskeleton influences Hippo pathway activity in R8 cells is currently unclear, but seems to be distinct from those that operate during organ growth.

The Hippo pathway is important for cell fate determination in a number of tissues in addition to the R8 cells of the *D. melanogaster* eye, including the posterior follicle cells of the *D. melanogaster* egg chamber [90–93], the peripodial epithelium/disc proper cell fate decision of the *D. melanogaster* larval eye imaginal disc [94] and the inner cell mass/trophectoderm decision in the early mouse blastocyst [95–99]. In posterior follicle cells, as in R8 cells, some Hippo pathway components, such as the Fat branch of the pathway, are not involved in inducing cell fate [90–92]. Interestingly, mutations in *α-Spec* and *β-Spec*, but not in *kst* or *crb*, lead to overproliferation of posterior follicle cells and the upregulation of Yki target genes [53–55], indicating that the basolateral spectrin cytoskeleton, rather than the apical spectrin cytoskeleton, regulates the Hippo pathway in these cells. Combined with our results, this suggest that different cells have repurposed different components of the Hippo pathway to control cell fate. By defining the signalling logic employed by the Hippo pathway to control different cell fate choices and tissue growth, we should reveal new insights into these biological processes and also define how cellular machinery is redeployed in living systems.

## MATERIALS AND METHODS

### *Drosophila melanogaster* genetics

The following *D. melanogaster* stocks were used, many available from the Bloomington *Drosophila* Stock Centre (BDSC), the Vienna *Drosophila* Resource Centre (VDRC), the Kyoto Stock Centre (KSC) and the National Institute of Genetics in Japan (NIG): *lGMR-Gal4* (Claude Desplan), *UAS-β-gal RNAi*^*GD*^ (VDRC, #51446), *UAS-yki RNAi*^*KK*^ (VDRC, #104523), *UAS-wts RNAi*^*KK*^ (VDRC, #106174), *UAS-α-Spec RNAi*^*GD*^ (VDRC, #25387), *UAS-α-Spec RNAi*^*TRiP*^ (VDRC, #56932), *UAS-β-Spec RNAi*^*TRiP*^ (BDSC, #38533), *UAS-β-Spec RNAi*^*GD*^ (VDRC, #42053), *UAS-β-Spec RNAi*^*GD*^ (VDRC, #42054), *UAS-kst RNAi*^*TRiP*^ (BDSC, #33933), *UAS-kst RNAi*^*GD*^ (VDRC, #37074), *UAS-jub RNAi*^*GD*^ (VDRC, #38443), *UAS-jub RNAi*^*KK*^ (VDRC, #101993), *FRT82B crb*^*11A22*^ [71,100], *FRT82B crb^ΔFBM^ (Y10AP12AE16A).HA[w+ GMR]* [42], *UAS-crb*^*extra*^, *UAS-crb*^*intra*^ [101], *UAS-crb*^*intraΔFBM (Y10AE16A)*^, *UAS-crb*^*intraΔPBM (ΔERLI)*^ [102], *FRT82B kibra*^*4*^ [37], *Kst-Venus* (KSC, #115285), *Kibra-Venus*, *Mer-Venus* (to be described elsewhere).

*D. melanogaster* were raised at room temperature (22-23°C) or 18°C on food made with yeast, glucose, agar and polenta. Animals were fed in excess food availability to ensure that nutritional availability was not limiting. All experiments were carried out at 25°C. Males and females were used for all experiments. Mutant clones were generated using the *eyFlp/FRT* system to generate mutant clones in *D. melanogaster* eye [70].

### Immunostaining and microscopy

Dissections were performed as described in Hsiao, et al. [103]. Briefly, retinas were dissected in PBS and fixed in 4% paraformaldehyde, washed in PBS for one hour and rinsed in PBST (PBS with 0.3% Triton X-100). During this wash, the lamina was removed. Retinas with strong pigment were washed in PBST for 4-5 days, with the media refreshed once a day, to remove the pigment. Retinas were blocked in blocking solution (5% NGS in PBST) and incubated in primary antibody overnight. Following a one hour wash in PBST, retinas were incubated overnight in secondary antibody. Tissues were mounted in VectaShield Mounting Medium (Vector Laboratories, H-1000) or 90% glycerol on bridge slides [104]. The following primary antibodies were used: mouse anti-Rh5 (1:200, Claude Desplan), rabbit anti-Rh6 (1:1000, Claude Desplan), chicken anti-GFP (1:1000, Abcam, ab13970), rat anti-Ecad (1:50, Developmental Studies Hybridoma Bank (DSHB), DCAD2), rat anti-HA (1:100, Santa Cruz, 3F10), mouse anti-α-Spec (1:100, DSHB, 3A9), mouse anti-pSqh (1:50, Cell Signalling, 3671S). Secondary antibodies conjugated to Alexa405, Alexa488, Alexa555 and Alexa647 (Life Technologies and Invitrogen) were used at a concentration of 1:500. DAPI (1:500, Sigma-Aldrich, D9542) and Phalloidin-TRITC (1:200-500, Sigma-Aldrich, P1951) staining was completed before mounting. Images were collected on a Nikon C2 or Olympus FV3000 confocal microscope, or an Olympus FVMPE-RS multiphoton microscope.

### Image analysis and statistics

Images were analysed using FIJI/ImageJ (https://imagej.net/Fiji). All statistical analyses were completed in RStudio using the stats package. All graphs were generated in RStudio using the ggplot [105] and ggbeeswarm [106] packages. The number of R8 cells that expressed Rh5, Rh6 or both, were counted using the FIJI Cell Counter plugin. Retinas were scored only if there were more than 100 ommatidia in a single focal plane. Statistical comparisons between ratios of R8 subtypes was calculated from the total number of Rh5-positive cells (cells expressing only Rh5 and cells expressing both Rh5 and Rh6) using a two-tailed, unpaired t-test, with the following symbols used for p-value cut-offs: *** < 0.0001, ** < 0.001, * < 0.01, ns > 0.01. Error bars represent the standard deviation. Statistical comparison of the ratio of R8 subtypes in clonal tissues was calculated using an ANOVA and multiple comparisons between genotypes calculated using Tukey’s Honest Significant Difference test, with the following symbols used for p-value cut-offs: **** < 0.0001, *** < 0.001, ** < 0.01, * < 0.05, ns > 0.05.

## ACKNOWLEDGEMENTS

We thank C. Desplan for comments on the manuscript and reagents, and R. Johnston, H. Richardson, the Bloomington *Drosophila* Stock Center, the Vienna *Drosophila* RNAi Center, the Kyoto Stock Center, the Australian *Drosophila* Research Support Facility (www.ozdros.com), and the Developmental Studies Hybridoma Bank for *D. melanogaster* stocks and antibodies. K.F.H is a National Health and Medical Research Council Senior Research Fellow (APP1078220) and J.M.P. was supported by an Australian Postgraduate Award. This research was supported by the Australian Research Council (DP180102044). We acknowledge the Peter Mac Centre for Advanced Histology and Microscopy and support to them from the Peter MacCallum Cancer Foundation and the Australian Cancer Research Foundation.

---

**Figure S1.**
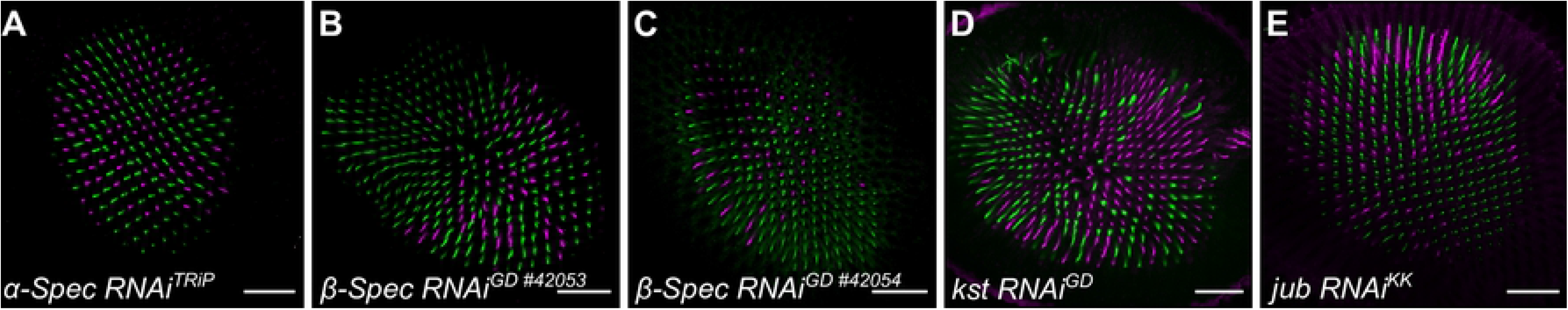
RNAi lines for Spectrins and *jub*. (**A-E**) Confocal microscope images of adult *D. melanogaster* retinas stained with anti-Rh5 (magenta) and anti-Rh6 (green) antibodies. All lines were driven by *lGMR-Gal4*. Retinas expressed *α-Spec RNAi*^*TRiP*^ (**A**), *β-Spec RNAi*^*GD* #42053^ (**B**), *β-Spec RNAi*^*GD* #42054^ (**C**), *kst RNAi*^*GD*^ (**D**) and *jub RNAi*^*KK*^ (**E**). Scale bars are 50μm.

**Figure S2.**
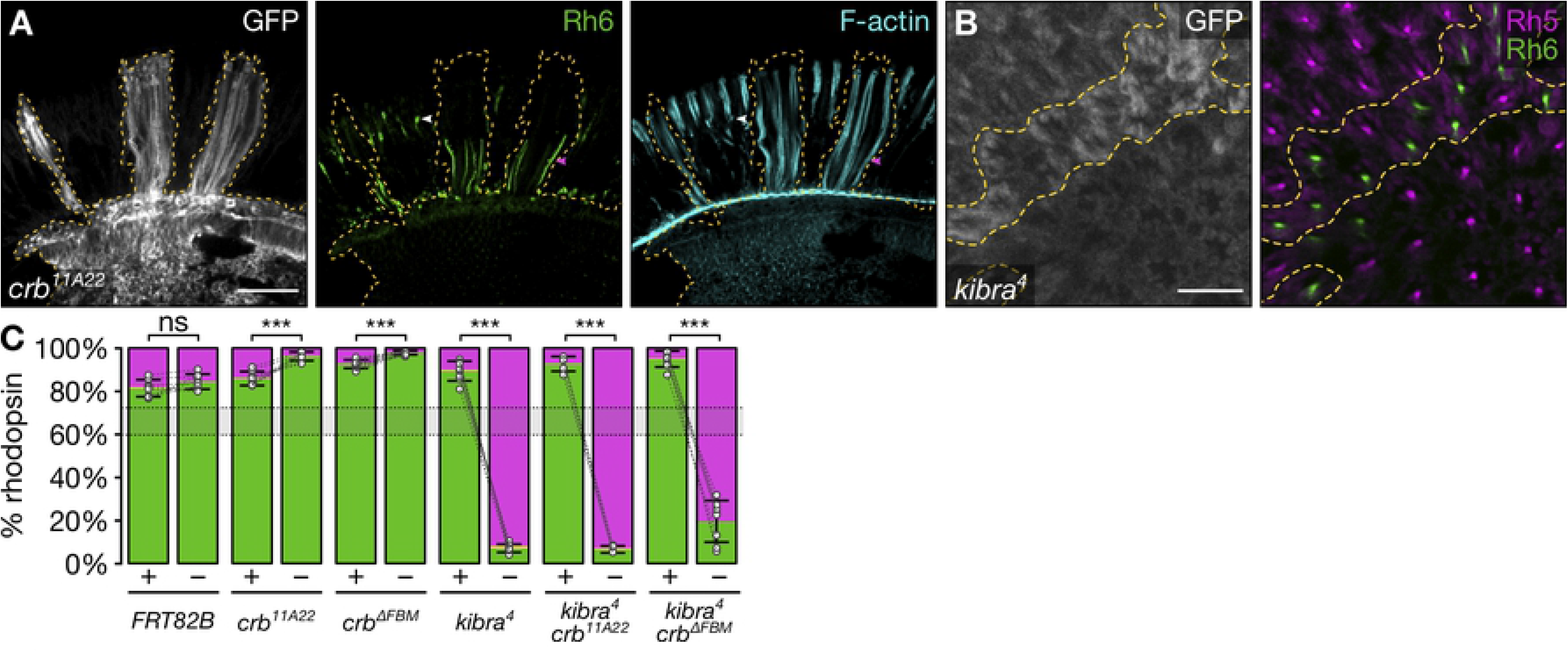
R8 cells harbouring a null c*rumbs* allele display rhabdomere morphology defects. (**A**) Confocal microscope images of adult *D. melanogaster* retinas stained with anti-GFP (grey) and anti-Rh6 (green) antibodies and stained with Phalloidin (cyan) to visualise the rhabdomere. GFP-negative clones were mutant for *crb*^*11A22*^. Arrowheads indicate a wild type ommatidium (magenta) and a mutant ommatidium (white). Scale bar is 50μm. (**B**) Confocal microscope images of adult *D. melanogaster* retinas stained with anti-GFP (grey), anti-Rh5 (magenta) and anti-Rh6 (green) antibodies. GFP-negative clones were mutant for *kibra^4^*. Scale bar is 50μm. (**C**) Proportion of R8 cells in wild type (‘+’) or mutant (‘−’) clones that express Rh5 (magenta), Rh6 (green), or both (yellow). Grey lines connect wild type and mutant clones from the same retina. The error bars represent the standard deviation of total % Rh5 (% Rh5 + % Rh5+Rh6). Total % Rh5 was compared with two-sided, unpaired t-test; ns = not significant, *** = p<0.0001. The shaded grey region between the dotted grey lines indicates wild type Rh5:Rh6 ratio range. *FRT82B*: n = 8 retinas, 4065 ommatidia; *crb*^*11A22*^: n = 8, 1394; *crb*^*ΔFBM*^.*HA*: n = 10, 3851; *kibra*^*4*^: n = 8, 2776; *kibra*^*4*^ *crb*^*11A22*^: n = 5, 1174; and *kibra*^*4*^ *crb*^*ΔFBM*^.*HA*: n = 8, 2479.

